# Boosting Live Malaria Vaccine with Cytomegalovirus Vector Can Prolong Immunity through Innate and Adaptive Mechanisms

**DOI:** 10.1101/2023.05.02.539025

**Authors:** Komi Gbedande, Samad A. Ibitokou, Monique L. Ong, Mariapia A. Degli-Esposti, Michael G. Brown, Robin Stephens

## Abstract

Vaccines to persistent parasite infections have been challenging, and current iterations lack long-term protection. Cytomegalovirus (CMV) chronic vaccine vectors drive protection against SIV, tuberculosis and liver-stage malaria correlated with antigen-specific CD8 T cells with a Tem phenotype. This phenotype is likely driven by a combination of antigen-specific and innate adjuvanting effects of the vector, though these mechanisms are less well understood. Sterilizing immunity from live *Plasmodium chabaudi* vaccination lasts less than 200 days. While *P. chabaudi*-specific antibody levels remain stable after vaccination, the decay of parasite-specific T cells correlates with loss of challenge protection. Therefore, we enlisted murine CMV as a booster strategy to prolong T cell responses against malaria. To study induced T cell responses, we included *P. chabaudi* MSP-1 epitope B5 (MCMV-B5). We found that MCMV vector alone significantly protected against a challenge *P. chabaudi* infection 40-60 days later, and that MCMV-B5 was able to make B5-specific Teff, in addition to previously-reported Tem, that survive to the challenge timepoint. Used as a booster, MCMV-B5 prolonged protection from heterologous infection beyond day 200, and increased B5 TCR Tg T cell numbers, including both a highly-differentiated Tem phenotype and Teff, both previously reported to protect. B5 epitope expression was responsible for maintenance of Th1 and Tfh B5 T cells. In addition, the MCMV vector had adjuvant properties, contributing non-specifically through prolonged stimulation of IFN-γ. *In vivo* neutralization of IFN-γ, but not IL-12 and IL-18, late in the course of MCMV, led to loss of the adjuvant effect. Mechanistically, sustained IFN-γ from MCMV increased CD8α^+^ dendritic cell numbers, and led to increased IL-12 production upon *Plasmodium* challenge. In addition, neutralization of IFN-γ before challenge reduced the polyclonal Teff response to challenge. Our findings suggest that, as protective epitopes are defined, an MCMV vectored booster can prolong protection through the innate effects of IFN-γ.

**Authors’ summary:** Malaria represents a challenging target for vaccination. This is in part because of the requirement for CD4 T cell immunity in addition to the standard B cell responses current vaccines induce. However, human malaria vaccine approaches thus far have limited longevity of protection due to decay of T cell responses. This includes the most advanced malaria vaccine, a virus-like particle expressing one recombinant liver-stage antigen (RTS,S), and liver-stage parasites attenuated by radiation (PfSPZ), as well as live vaccination with drug-treatment. Our work seeks to prolong this protection using MCMV, a promising vaccine vector known to promote CD8 T cell responses. We observed that boosting the live malaria vaccine with MCMV including a *Plasmodium* antigen led to longer protection from *P. chabaudi* parasitemia, and can be used to promote maintenance of antigen-specific CD4 T cells. In investigating the mechanisms of the MCMV booster, we found that the cytokine IFN-γ is required for prolonged protection and enhances priming of the innate immune system for prolonged protection from malaria. Our research informs both the quest for a longer-lived malaria vaccine and that to understand mechanisms of protection from persistent infection.

## Introduction

Compared to acute and localized infections, persistent systemic infections are the most challenging vaccines to develop [1, 2]. Despite impressive progress for malaria vaccine development, and using emerging approaches including live or attenuated parasites, and recombinant protein constructs targeting different stages in the parasite life cycle, none of the current vaccine candidates provides both highly effective and long-lived immunity. Both natural parasite infection and malaria vaccines generate immunity to parasitemia on re-infection that wanes quickly, and this is shown to be due to a progressive loss of specific T cells [3, 4]. Key insights to decay of protection from persistent systemic infections has come from the study of immunity to parasite infections with *Leishmania* and *Plasmodium spp.,* which decays over time in the absence of infection [3, 5, 6]. Protection from recurrent infection in nature is likely mediated by sustained generation of otherwise short-lived Th1 Teff while infection persists [7, 8]. We and others have shown that both short-lived effector T cells and long-lived effector memory T cells are generated in persistent infection. It is unclear if long-lived memory T cells (Tmem) play an important role, or could be used to improve protection from potentially persistent systemic infections. Persistent parasite infection, including *Leishmania major* or *P. chabaudi,* generate recirculating Tcm that do not protect as well as activated Teff or Tem, defined as previously divided CD4^+^CD62L^lo^ Tem/Teff [9], or distinguished as CD127^-^ Teff and CD127^hi^ CD44^hi^ CD62L^lo^ CD27^-^ Tem [10]. In *P. chabaudi* infection, we showed that inducing a pre-activated MSP1-specific B5 T cell receptor transgenic (TCR Tg) T cell response, along with a B cell response to MSP-1 p19, before infection with *P. chabaudi*, was an effective way to promote an effective reduction of the parasitemia of re-infection. These data suggest an important role for effector T cells in protection. It should be noted that while B5 T cells protect immunocompromised recipients from death, they do not eliminate parasite in the absence of B cells [11]. By defining Teff as CD127^-^, an early marker of T cell activation that is transient and not present on Tem, we were able to determine that adoptively transferred Teff are the most protective T cells in systemic, blood-stage *P. chabaudi* infection [8, 10]. However, short-lived CD4 T cell immunity has been identified as the crucial variable resulting in short-term protection from persistent *P. chabaudi* infection or vaccination [3, 12]. However, standard vaccines do not generate durable Teff responses, leading to the problem at hand of what vaccine vector to use to prolong protection, if the cell type that protects is short-lived.

Therefore, we investigated the potential of CMV to increase the duration of the T cell response to effectively prolong vaccine-induced immunity to *Plasmodium* infection as a booster vector. CMV has emerged as a promising long-lasting vaccine vector, for example, it provided lasting immunity over a year against mucosal Simian Immunodeficiency virus (SIV) challenge in Rhesus macaques [13]. CMV is a potent CD8 T cell-inducing vector with efficacy against SIV, ebolavirus, tuberculosis, cancer and liver-stage malaria [14–18]. CMV infection can induce sustained, or inflationary CD8 T cell responses with an effector memory phenotype. Th1-type effector function is also generated [19–21], but the effect of persistent CMV on CD4 T cells is less well understood. CMV also has non-specific effects suggesting a useful adjuvant effect of the vector. For example, MCMV provides non-specific protection to *Listeria monocytogenes*, *Yersinia pestis* and influenza virus, particularly in the latent phase of infection [22, 23]. Previous vaccine studies all have used CMV in the priming phase of the vaccine response, and it has not been tested as a vector for boosting existing responses, nor has it been studied extensively in the generation of CD4 T cell responses. In addition, CMV generates effector/memory T cells, but the specific phenotype has not been determined, and extending the duration of an effector response will be critical to the fight against persistent infection.

In this study, we found that the MCMV vector boosted protection generated by a live malaria vaccine against challenge infection with heterologous parasite. In addition to showing that we can induce epitiope-specific effector T cells 40-60 days post-boost, we identified a significant adjuvanting effect of boosting with MCMV. We showed that IFN-γ produced during the persistent phase of MCMV increases CD8α^+^ dendritic cell numbers. Upon challenge, MCMV boosted animals had increased serum IL-12 and a larger CD4 polyclonal effector T cell response, which were reduced by neutralization of IFN-γ before infection. This work informs our basic understanding of the immunology and vaccinology of persistent infections, and suggests that persistent stimulation of innate and adaptive mechanisms can promote persistence of protective immunity.

## Results

### MCMV as a vaccine vector with adjuvant properties against *P. chabaudi*

In order to evaluate an MCMV vaccine vector and observe vector- and *Plasmodium*-specific CD4 T cell phenotypes, we generated an MCMV vector carrying the *P. chabaudi* MSP-1 B5 CD4 T cell epitope (MCMV-B5) using bacterial artificial chromosome (BAC) recombineering. The B5 epitope was inserted in the intermediate early 1 locus with a linker to facilitate processing, as diagrammed in **S1A Fig**. As MCMV has been shown to induce more sustained CD4 T cell numbers in BALB/c, than in C57BL/6 mice [24], we developed a BALB/c system. Although *P. chabaudi* MSP-1 B5 CD4^+^ T cells prevent immunodeficient animals from dying, B5 T cells alone do not affect *P. chabaudi* parasitemia significantly unless they are pre-activated, and in the presence of parasite-specific B cells [11]. Several aspects of the virology and immunology of the two constructs were compared for rigor of the study. The kinetics and tissue tropism of MCMV-B5 infection was confirmed to be the same as the parental virus, MCMV-BAC using plaque assay (**S1B Fig**) and PCR (**S1C Fig**). The virus is largely cleared by day 30, however low recrudescence can be seen at later timepoints. Proportions of polyclonal CD4^+^CD127^-^ Teff, and CD4^+^ CD44^hi^CD127^hi^ Tmem were similar for MCMV-BAC and MCMV-B5 at day 15 p.i. (**S1D Fig)**. MCMV-specific CD4^+^ T cells were identified using m78 peptide/I-A^d^ tetramers, using CLIP/I-A^d^ as non-specific tetramer control, and MCMV-specific CD8^+^ T cells were detected with the m45/D^d^-tetramer [20, 21]. Both MCMV-BAC and MCMV-B5 induced an MCMV m78-specific CD4 T cell response at day 15 p.i. (**S1E Fig**). A large MCMV m45-specific CD8 T cell response to MCMV-B5 was sustained near a million T cells per spleen from day 11 to 60 p.i. (**S1F Fig**).

MCMV is an excellent vaccine vector that showed early promise against liver-stage malaria [16] and other infections fought by CD8 T cells. However, the mechanisms by which the vector contributes to the adjuvanting protective effect are not well understood. To test for a protective effect of the low chronic phase of MCMV infection on *P. chabaudi* challenge, BALB/cByJ mice were infected with MCMV-B5 or MCMV-BAC or aged, and challenged 60 days later with *P. chabaudi* AS, as shown schematically in **Fig 1A**. Groups given either MCMV-BAC or MCMV-B5 had strongly reduced parasite growth upon *P. chabaudi* infection compared to a primary infection age-matched control (**Fig 1B**). Pathology from malaria, as indicated by weight loss, was also significantly inhibited by prior MCMV-BAC or MCMV-B5 infection (**Fig 1C**). This strong, non-specific effect reduced parasitemia and pathology by about half, so not as effectively as prior infection, or live vaccination [3].

**Figure 1.**
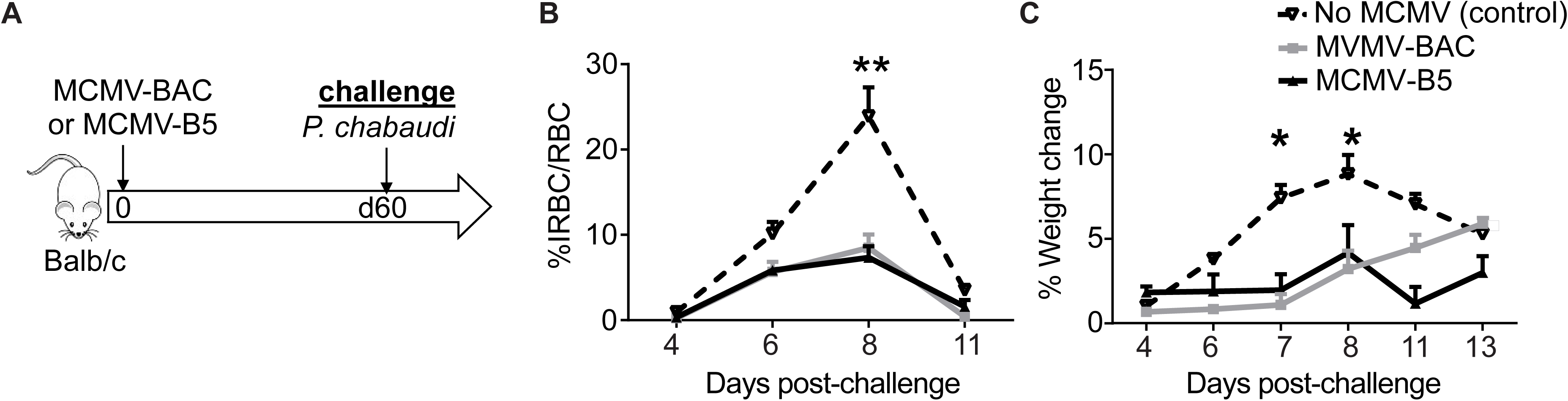
Prior MCMV infection protects against *P. chabaudi* infection. **(A)** Experimental schematic shows mice infected with MCMV-B5 or -BAC and challenged on day 60 post MCMV infection with *P. chabaudi* AS. **(B)** Parasitemia and **(C)** weight loss were determined after challenge. Data are representative of three independent experiments with 5 animals per group. Groups were compared using ANOVA followed by Tukey’s post-test with *, *P* < 0.05; **, *P* < 0.01. Mean shown with error bars representing SEM.

### MCMV stimulates and maintains *Plasmodium*-specific T cell responses

Using B5 TCR Tg T cells, we were able to distinguish the non-specific adjuvant effects from epitope-specific effects on CD4^+^ T cells [11]. To determine effects of the MCMV vector on epitope-specific T cells, we adoptively transferred B5 Tg T cells (2 x 10^6^) and infected the recipients with MCMV-BAC or MCMV-B5, as illustrated in **Fig 2A**. The B5 CD4 T cell response in the spleen 15 and 55 days post-MCMV infection was assessed by flow cytometry. The proliferated B5 T cell (CTV^-^ Thy1.2^+^) number was specifically increased by MCMV-B5, which leading to a specific increase that lasted until day 55 (**Fig 2B**). MCMV-BAC infected mice also maintained proliferated CTV^-^ B5 T cell numbers slightly above background to day 55. In order to test the MCMV-driven generation of effector T cells, they were stained for the IL-7Rα (CD127), which is downregulated on recently activated T cells. The majority of CTV^-^ B5 TCR Tg T cells on day 15 are CD127^-^ Teff (**Fig 2C**). On day 55, the fraction of Tmem increases, but a significant number of Teff also are present, as clearly seen in the flow cytometry plot of B5 T cells recovered at this timepoint (**Fig 2D**). In addition, we observed that MCMV-B5, but not MCMV-BAC, specifically promoted the differentiation of MCMV-B5 epitope-specific GC Tfh (CXCR5^hi^PD-1^hi^, **Fig 2E**).

**Figure 2.**
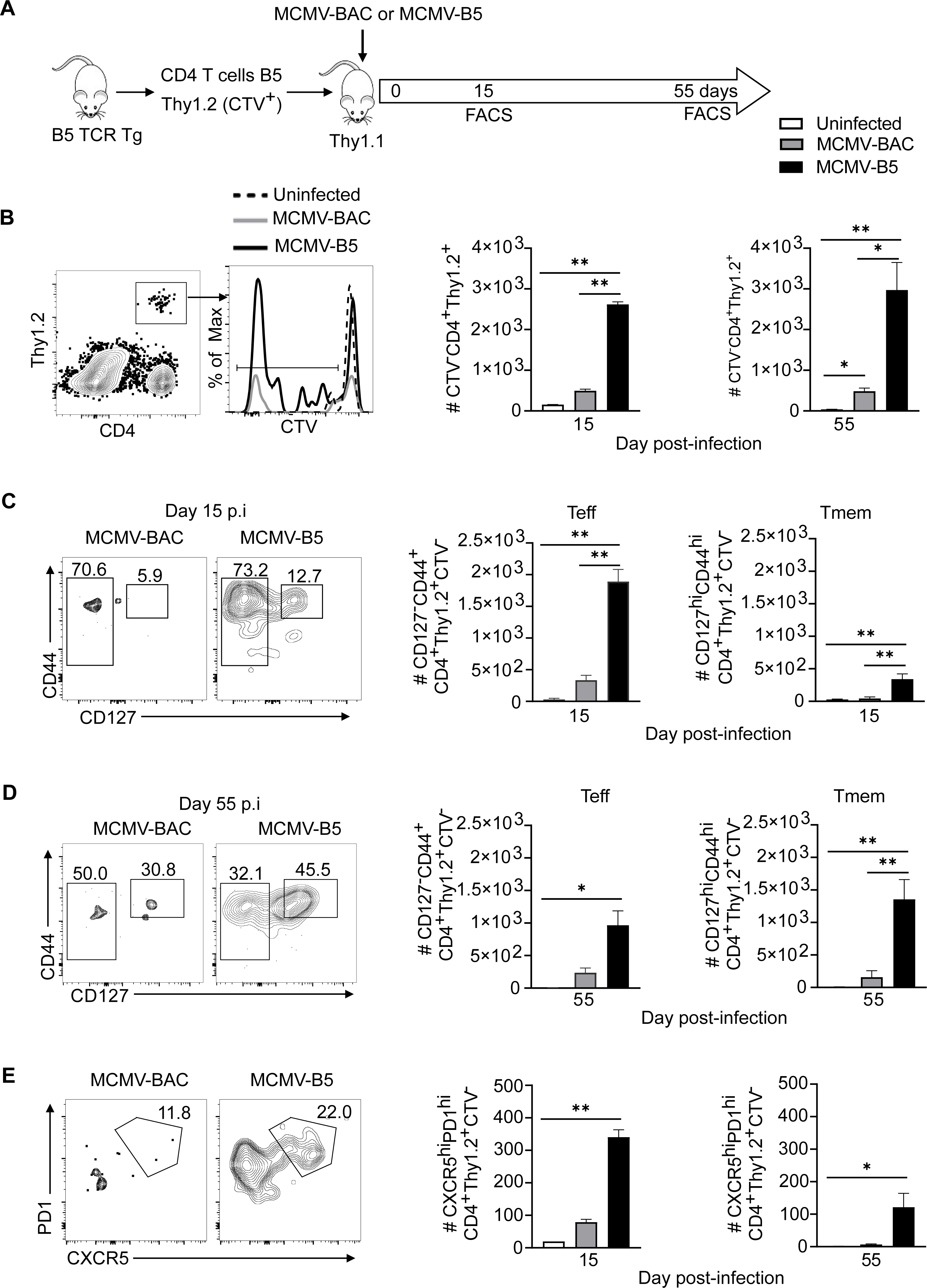
MCMV-B5 stimulates and maintains parasite-specific effector T cell responses. **(A)** Experimental schematic shows MSP-1 specific B5 TCR Tg CD4 T cells (CTV^-^Thy1.2^+^) were adoptively transferred into Thy1.1 mice that were then infected with MCMV-BAC or MCMV-B5. Splenic T cells were gated on B5 TCR Tg (CD4^+^CTV^-^Thy1.2^+^) recovered at days 15 and 55 post-MCMV infection. **(B)** Plot, histogram and graphs show gating of proliferated B5 T cells. **(C)** Plots and graphs show B5 Teff (CD4^+^Thy1.2^+^CTV^-^, CD127**^-^**CD44^hi^) and Tmem (CD4^+^Thy1.2^+^CTV^-^, CD127^hi^CD44^hi^) **(D)** on day 15, or **(E)** day 55. Plots and graphs show B5 GC Tfh days 15 and 55 p.i.. Plots are concatenated from 3-5 animals per group. Data are representative of two experiments with 3-5 mice in each group. Mean shown with error bars representing SEM and analyzed using One-Way ANOVA followed by Tukey’s post-test; *, *P* < 0.05; **, *P* < 0.01.

We also detected some MCMV-specific CD4 T cell responses using MCMV m78 tetramer at 11 and 60 days after MCMV-B5 infection (S2A **Fig**). MCMV-B5 infection induced MCMV m78-specific CD4^+^ T cell responses that persisted for two months (S2B **Fig**). Phenotypically, MCMV vector-specific T cells were predominantly Teff (Tet^+^ CD127^-^) early in infection (day 11), with re-upregulation of CD127 into a Tmem phenotype at day 60 (S2C **Fig**), similar to the B5-specific response to MCMV-B5. Proliferation of transiently transferred B5 T cells in response to MCMV-B5 was tested over various five-day windows. Congenic recipients (Thy1.1) were infected with MCMV-BAC or MCMV-B5 or uninfected, and CTV-labelled CD4-purified B5 TCR Tg T cells were adoptively transferred for the last five days of infection, testing presentation of B5 peptide between days 5-10, 25-30, or 55-60 of MCMV infection. Splenocytes were analyzed for proliferation (%CTV-) of B5 T cells in each experimental window (**S2D Fig**). The MCMV-B5 vector, and not the MCMV-BAC control, led to strong specific proliferation of epitope-specific B5 Tg T cells during acute infection (days 5-10) (**S2E Fig**). Despite the presence of B5 TCR Tg Teff on day 55, there was no detectable proliferation in this assay at later timepoints. This may be due to background proliferation, even in uninfected animals.

### MCMV boosts long-term protection of live malaria vaccine

As the protection of malaria vaccines tested in mice and humans decays after one to two years [25, 26], we sought to test a vaccine booster strategy to prolong immunity. *Plasmodium spp*. infection followed by drug treatment is an effective vaccination protocol in C57BL/6 mice, and also in human studies; however, this protection shows measurable decay by day 200 in mice [3, 27]. Using the protocol published by Freitas do Rosario et al, where mice are infected with *P. chabaudi* AS followed by sub-curative dose of chloroquine each time parasitemia reaches 1%, we tested sterilizing immunity with heterologous reinfection (**S3A Fig**). This experiment confirmed that *P. chabaudi* infection induced sterilizing immunity for two months, and that a measurable decay in protection was evident by day 200 in BALB/c mice (**S3B Fig**), as in B6. We also confirmed that there was not a measurable decay in antibody concentration between 2 and 6.5 months p.i., as shown previously in B6 mice (**S3C Fig**), leading us to test a known T cell-response-promoting vector to attempt to prolong T cell responses and protection through as yet unknown adjuvant effects.

To test the potential of an MCMV-based booster vaccine, mice were vaccinated with live *P. chabaudi* AS infection followed by sub-curative doses of chloroquine and then boosted with MCMV-BAC or MCMV-B5 140 days later, as schematically depicted in **Fig 3A**. As vaccine-induced protection decays by day 200, we tested MCMV boosters for protection from heterologous re-infection with *P. chabaudi* AJ at day 200, which corresponds to 60 days after boosting. This timing was chosen on the basis that neither immunity nor the T cell response to live vaccination has completely decayed by day 120 [3, 12], and that the MCMV infection is below detectable by day 60 [24], though we expected it to show prolonged stimulatory capacity at this timepoint [22]. Age- and sex-matched *Plasmodium-*vaccinated mice, with or without the MCMV-booster, were challenged with heterologous *Plasmodium* infection. The booster prolonged live vaccine induced protection, as evidenced by the data shown in **Fig 3B** that mice that were boosted with MCMV had a lower parasitemia on challenge on day 200 post-vaccination (p.v., average of 0.06%, or 0.05% without or with the B5 epitope, respectively), on day 4 post-challenge (p.c.), than vaccinated mice challenged without boosting (1.57% on day 4 p.c., and 0.5% on day 6 p.c.). The boosting effect does not require the B5 epitope in a significant way, which is not surprising, B5 CD4 T cell epitope was included as a way to track specific T cells, not to protect as a single epitope. Heterologous parasite-specific serum antibody was measured by ELISA at day 7 p.c. (**Fig 3C**). As previously reported parasite-specific IgG responses remain stable up to day 200 post-live vaccination [3], ELISA showed that levels of *P. chabaudi* AJ-specific serum IgG in the first week of challenge were not altered by the MCMV booster (**Fig 3C**). The boost was specifically chosen to promote T cell responses to support the ongoing antibody response generated by the live vaccine. In summary, *Plasmodium* vaccination efficacy, but not serum antibody, fades with time, and boosting with MCMV prolonged protection.

**Fig 3.**
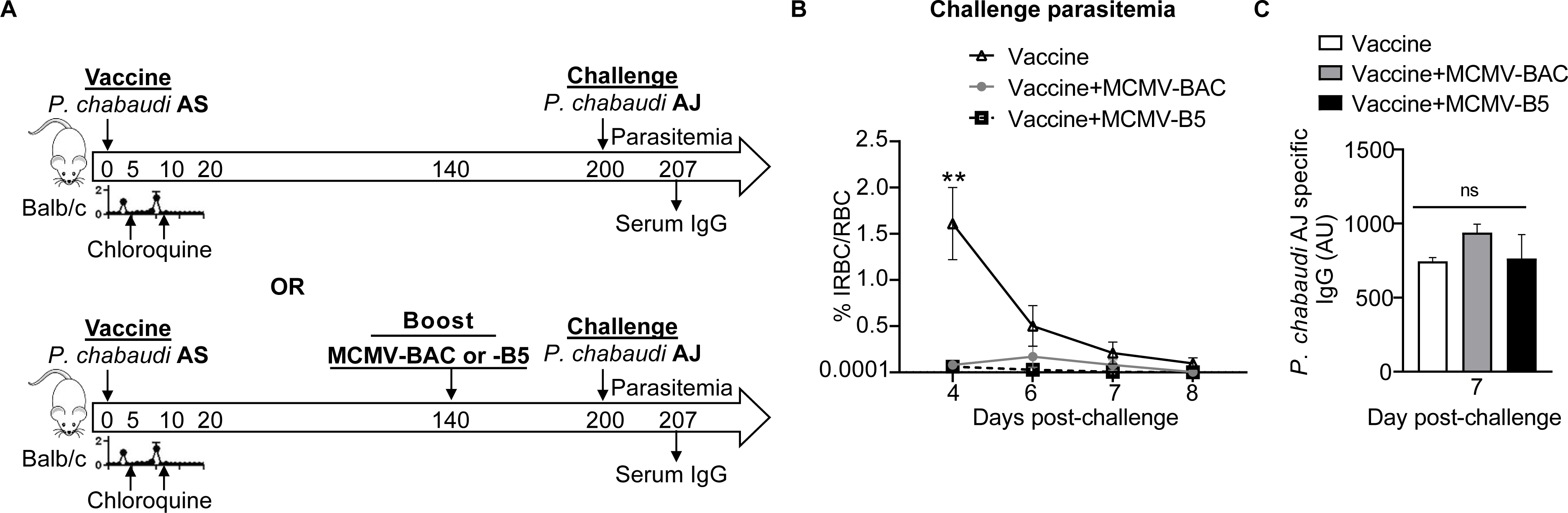
MCMV boosts long-term protection of live malaria vaccine. **(A)** Experimental schematic shows infection and drug-cure live vaccine combined with MCMV boosting strategy for protection to heterologous rechallenge. Mice were infected with *P. chabaudi* AS and treated with two subsequent non-curative doses of chloroquine (CQ). Some groups received MCMV-BAC or MCMV-B5 booster at day 140 (bottom timeline). All mice were challenged with heterologous *P. chabaudi* AJ strain 200 days after live vaccination. Graph showing **(B)** parasitemia of *P. chabaudi* AJ challengeand **(C)** serum level of *P. chabaudi* AJ-specific IgG antibody at day 7 post-challenge. Data are representative of two independent experiments with five mice per group showing mean, with error bars representing SEM and analyzed using ANOVA followed by Tukey’s post-test with **, *P* < 0.01; n.s., not significant.

### MCMV can boost parasite-specific T cell immunity induced by live malaria vaccine

CMV is a promising CD8 T cell-inducing vaccine vector and generates Tem and inflationary effector T cells [20, 28]. Less is known about CD4 T cell phenotypes in response to MCMV [20], and the potential for continuous generation of phenotypic effector CD4 T cells in the persistent phase is not well understood. In order to determine if MCMV-B5 boosted *Plasmodium*-specific CD4 T cell responses, B5 CD4 TCR Tg cells (Cell Trace Violet (CTV)-labelled) were adoptively transferred into congenic recipients (Thy1.1 BALB/cByJ), which were then immunized with live *P. chabaudi* vaccine, and boosted with MCMV-B5 or MCMV-BAC at day 140 p.v. (**Fig 4A**). B5 T cell effector and effector memory phenotype and cytokine responses were quantified at day 200 p.v.. Using MCMV-B5, as a booster to the live malaria vaccine specifically and significantly increased the numbers of divided B5 TCR Tg T cells (CD4^+^CTV^-^Thy1.2^+^) present on day 200 p.v., while MCMV-BAC did not (**Fig 4B**). Therefore, more transferred B5 T cells survived to day 200 after MCMV-B5 boost. B5 TCR Tg cells are not detectable after 200 days in the absence of parasite [8]. The MCMV-B5 boost 60 days earlier specifically increased the numbers of both CD127^-^ B5 Teff and CD127^hi^ B5 Tmem present at day 200, while MCMV-BAC did not increase B5 T cell numbers (**Fig 4C**). Among Tmem subsets, the number of both CD27^+^ and CD27^-^ effector memory T cells (CD44^hi^CD127^hi^CD62L^lo^, Tem) were significantly higher in MCMV-B5 boosted mice, while central memory (CD62L^hi^ Tcm) were not increased by MCMV boost (**Fig 4D**). In addition, the MCMV-B5 booster specifically generated or maintained functional B5 T helper type 1-type cells, as shown by their production of IFN-γ (**Fig 4E**), TNF (**Fig 4F**) and IL-2 (**Fig 4G**) at day 200. This data show that the vector generated well-presented epitopes, and could be used as a booster vaccine for T cells. In addition, the vector induced effector T cells for a longer time than reported for acute stimulation, in addition to making Tem.

**Fig 4.**
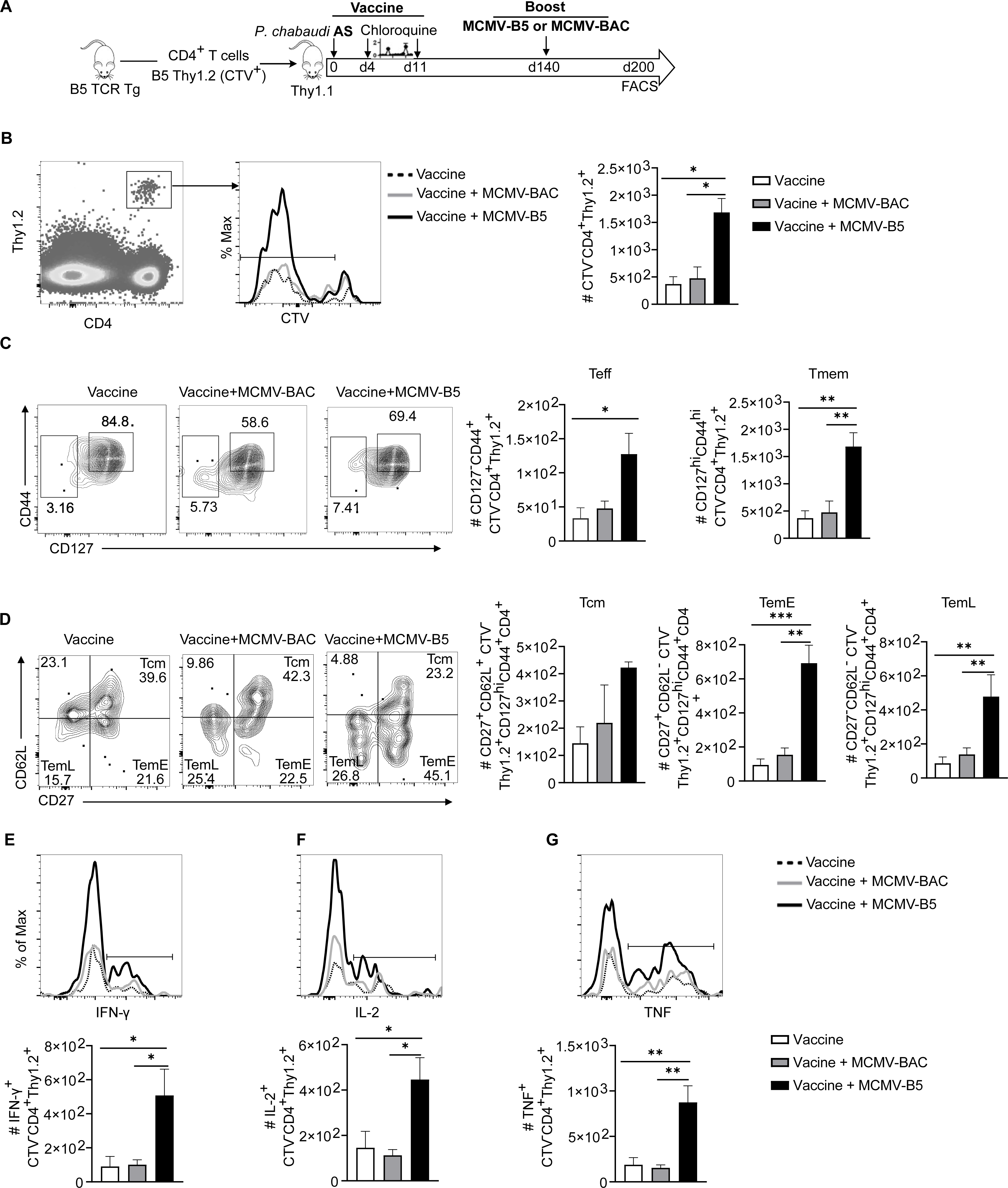
MCMV-B5 promotes maintenance of *Plasmodium*-specific effector T cells. **(A)** Experimental schematic shows live vaccine combined with MCMV boosting strategy with adoptively transferred CD4^+^ *Plasmodium* MSP1-specific T cells (B5 TCR Tg). CD4^+^ Thy1.2^+^ CTV^+^ B5 T cells were adoptively transferred into Thy1.1 mice that were then vaccinated. Recipient mice received MCMV-BAC or MCMV–B5 boosters at day 140, with T cell phenotyping at day 200 post-vaccination (p.v.). **(B)** Plot, histogram and graph show gating and quantification of parasite-specific CTV^-^ B5 T cells recovered on d200. **(C)** Plots and graphs showing number of Teff (CD127^-^) and Tmem (CD127^hi^CD44^hi^) out of CTV^-^ B5 TCR Tg cells on day 200. **(D)** Plots and graphs showing percentage and number of B5 memory T cell subsets gated on CD44^hi^CD127^hi^, Tcm (CD62L^+^CD27^+^), Tem^Early^ (TemE, CD62L^lo^CD27^+^) or Tem^Late^ (TemL, CD62L^lo^CD27^-^). Cytokines in B5 T cells were quantified and shown as histograms and graphs quantifying **(E)** IFN-γ, **(F)** IL-2- and **(G)** TNF-positive CTV^-^ B5 T cells. Data are representative of two independent experiments. Plots are concatenated from 5 animals per group, while graphs include individual animals and show mean with error bars representing SEM. Groups were analyzed using One-Way ANOVA followed by Tukey’s post-test with *, *P* < 0.05; **, *P* < 0.01, ***, *P* < 0.001.

### IFN-γ, but not IL-12 and IL-18, induced by MCMV, is associated with protection from *P. chabaudi* infection

As we had observed a protective effect by the MCMV-BAC, we took the opportunity to understand mechanisms explaining the observed benefits of using this virus as a vaccine vector in addition to the ability to present T cell epitopes and generate Teff. To determine if cytokines induced by long-term MCMV infection are mechanistically responsible for promoting protection from heterologous challenge, we used neutralizing antibodies to test the roles of IFN-γ, as well as the combination of IL-12 and IL-18, in the period before challenge after 40 days of MCMV infection. IFN-γ was neutralized starting on day 34 of MCMV-BAC infection, and before *P. chabaudi* challenge, as shown schematically in **Fig 5A**. *In vivo* neutralization of IFN-γ after MCMV, but before *P. chabaudi* infection was associated with a substantial loss of protection against *P. chabaudi* challenge. The mice treated with anti-IFN-γ before infection had significantly higher parasitemia compared to isotype-treated controls previously infected with MCMV (**Fig 5B**). The level of parasitemia in mice receiving neutralizing antibody is similar to a first infection suggesting that all or most of the reduction due to MCMV is due to IFN-γ. The effect of neutralizing MCMV-induced IFN-γ on challenge is also reflected in increased pathology, as measured by weight loss (**Fig 5C**), and hypothermia (**Fig 5D**). To determine if there is an effect of MCMV persistence-induced IFN-γ on the T cell response to challenge, the polyclonal CD4^+^ T cell phenotype was examined on day 15 post-challenge with *P. chabaudi.* Surprisingly, there was a significant reduction in responding CD4^+^ Teff after pre-treatment with anti-IFN-γ (**Fig 5E**).

**Fig 5.**
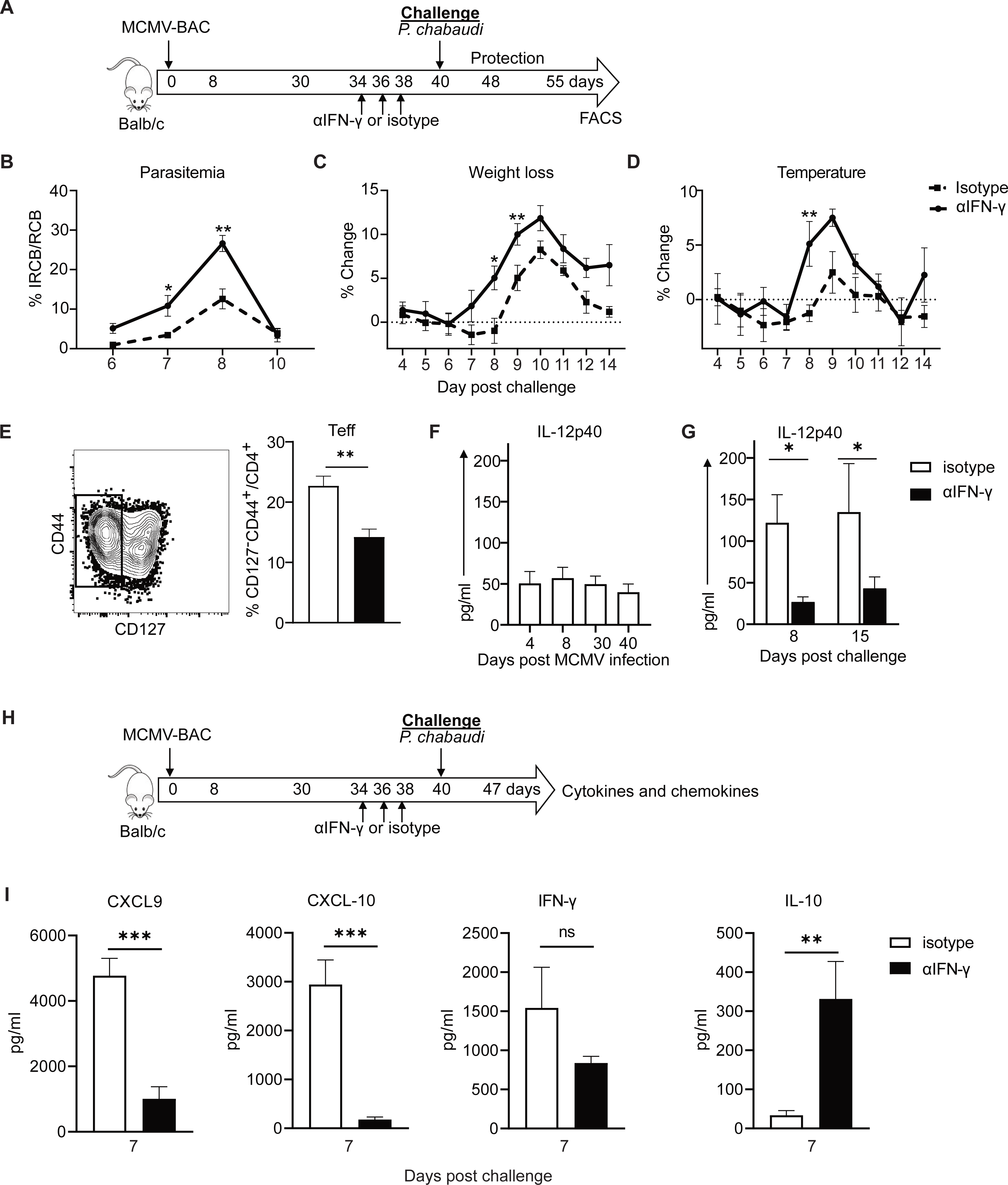
IFN-γ induced by MCMV leads to increased cytokines and protection from *P. chabaudi* infection. **(A)** Experimental schematic shows mice infected with MCMV-BAC and anti-IFN-γ or isotype control was administered at indicated days (34, 36 and 38 p.i.) prior to challenge with *P. chabaudi* at day 40. Graphs show **(B)** percentage of parasitemia, **(C)** weight loss and **(D)** hypothermia upon *P. chabaudi* challenge. **(E)** Plot and graph show fraction of polyclonal effector T cells at day 15 p.c., isotype in white and anti-IFN-γ in black bars. Graphs show plasma IL-12 p40 levels **(F)** throughout MCMV infection and **(G)** after *P. chabaudi* challenge. **(H)** Graphs show chemokines and cytokines on day 7 of *P. chabaudi* challenge after MCMV-BAC and antibody treatment. Graphs include 5 animals per group and show mean with error bars representing SEM. Multiple comparisons Student *t* test and Unpaired or Mann Whitney were used for all except (F) which was analyzed with One-Way ANOVA. *, *P* < 0.05; **, *P* < 0.01, ***, *P* < 0.001; n.s., not significant.

Evaluation of serum cytokines in MCMV infected animals showed IL-12p40 production throughout MCMV infection (**Fig 5F)** that is increased upon *P. chabaudi* challenge (**Fig 5G**). Strikingly, IL-12 p40 production on days 7, 8 and 15 p.c. as well as TNF, CXCL-9 and CXCL-10 on day 7 p.c. were significantly reduced by neutralization of IFN-γ for six days before challenge while IL-10 level was increased at 7 days p.c. (**Fig 5H)**. We also measured other cytokines including IL-6, IL-15, IL-18, IFN-α and IFN-β, which are known to be produced during MCMV infection to see if they were affected by neutralization of IFN-γ but did not see significant differences at these time points.

To investigate potential direct effects of neutralizing IFN-γ on challenge parasitemia due to persisting antibody and determine the role of persistent IFN-γ induced by MCMV in protection, we examined the effect of anti-IFN-γ clone H22 on *P. chabaudi* infection with or without an earlier dose of MCMV (**S4A Fig**). We found that administration of neutralizing antibody up to 2 days before infection with *P. chabaudi* does not have a significant measurable effect on peak parasitemia (**S4B Fig**), while it does affect parasitemia significantly, as above, if MCMV is given before infection (**S4C Fig**). To determine if neutralizing levels of antibody remained in the serum during challenge, we developed an assay to test induction of CXCL10, which is induced in macrophages by IFN-γ. CXCL10 mRNA was measured after culture of RAW 264.7 cells with serum from mice before challenge (d40, **S4D Fig**), and on day 7 post *P. chabaudi* challenge with or without prior anti-IFN-γ administration (**S4E Fig**). We found no detectable change for the relative CXCL10 mRNA level with Raw 264.7 cells cultured with d40 or d47 serum, though there was a trend suggesting antibody was there. Adding recombinant IFN-γ (rIFN-γ) to the assay allowed detection of neutralizing antibody. In this assay, a reduced level of CXCL10 mRNA was detectable in the presence of serum from 2 days after anti-IFN-γ (**S4F Fig**) and day 7 *P. chabaudi* challenge (**S4G Fig**). Therefore, though neutralizing antibody does remain in the system during *P. chabaudi* challenge, we conclude that the large effect of blocking IFN-γ after MCMV, and before challenge, is mostly due to inhibition of MCMV-induced not *P. chabaudi*-induced IFN-γ. Therefore, we investigated the mechanisms of this strong effect.

As IL-12 and IL-18 can induce IFN-γ from NK cells or previously stimulated T cells, we queried their role in induction of IFN-γ by MCMV, but did not find any effect of blocking IL-12 and IL-18 together in the time period before challenge on protection from *P. chabaudi* parasitemia or pathology by the MCMV vector (**S5 Fig**).

To explore the effect of IFN-γ on persisting epitope-specific T cells and the innate response in the context of late MCMV infection, we adoptively transferred CTV-labelled B5 TCR Tg T cells into Thy1.1 mice and then infected them with MCMV-B5, as illustrated in **Fig 6A**. IFN-γ was neutralized on days 44-48 of MCMV infection. Blocking IFN-γ induced by MCMV-B5 infection did not change the number of CTV^-^ B5 T cells at day 50 p.i. (**Fig 6B**). IFN-γ also did not affect the phenotype, as measured by the fraction of both Teff and Tmem (**Fig 6C**), or representation of Tmem subsets, as defined by CD27 and CD62L expression (**Fig 6D**). B5 T cells induced by MCMV-B5 are mostly memory T cells, with a fraction of effector T cells that is larger than that seen on day 60 of *P. chabaudi* infection [8]. A panel of innate cell markers showed that after IFN-γ neutralization on day 50 post-MCMV infection, the number of CD8α^-^ DCs (CD3^-^ CD11c^hi^MHCII^hi^) remained constant while the number of CD8α^+^ DCs showed a trend towards a reduction (**Fig 6E**). The level of IL-12p40 in the serum on day 50 post-MCMV infection was significantly reduced in the group that received IFN-γ neutralizing antibody compared to isotype (**Fig 6F**). Splenic CD8α^+^ DCs are the major producers of IL-12p40 in response to systemic stimuli [29].

**Fig 6:**
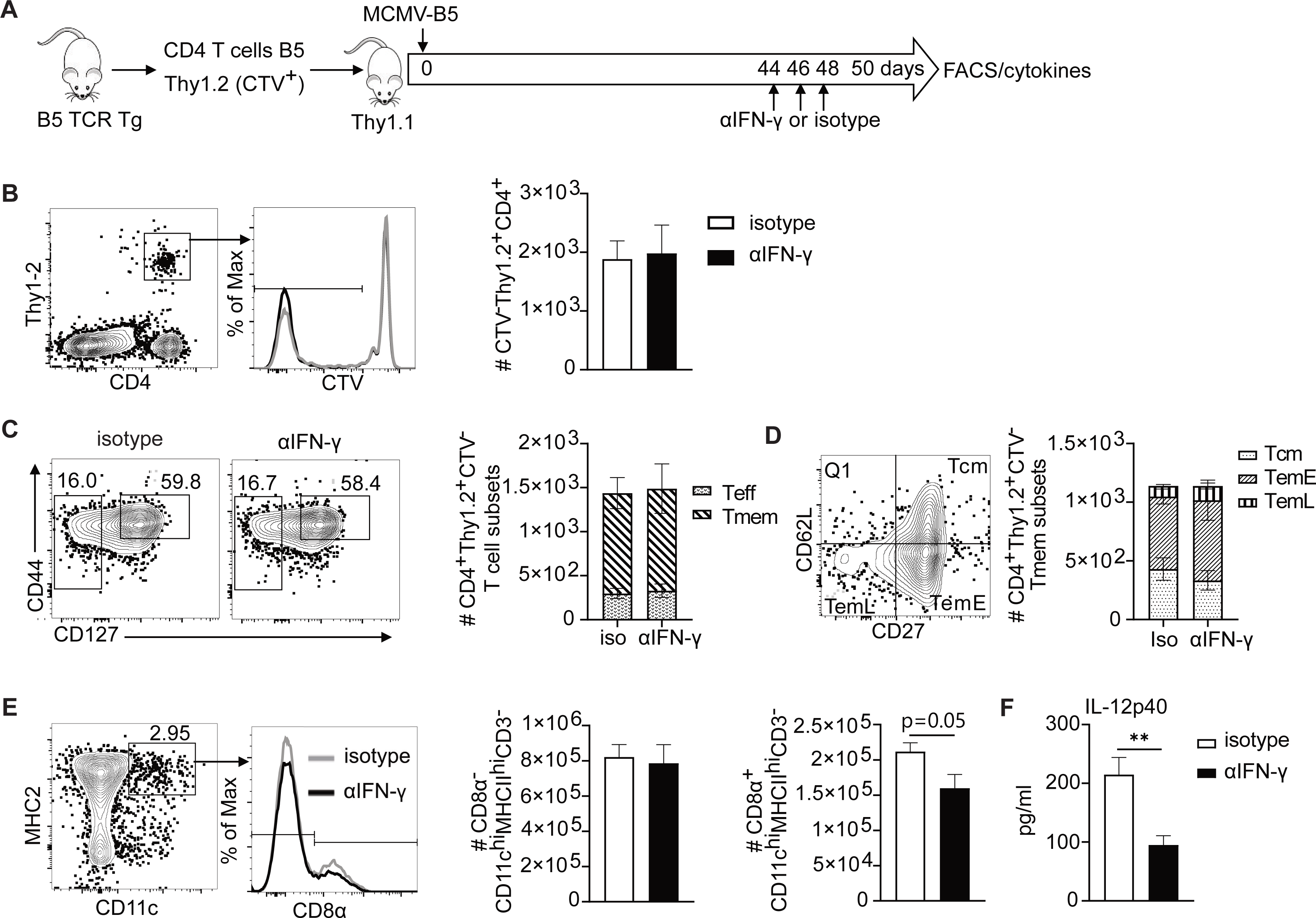
IFN-γ induced by MCMV does not regulate MCMV driven T cell phenotype, but does induce IL-12 p40. **(A)** Experimental schematic shows CTV-labelled B5 TCR Tg T cells were adoptively transfer into Thy1.1 (Balb/c) mice which were then infected with MCMV-B5 and received anti-IFN-γ antibody or isotype at indicated time points and spleen phenotyped for dendritic cells and B5 TCR Tg (Thy1.2^+^) CD4^+^ T cell at day 50 p.i.. **(B)** Plot, histogram and graph show gating and quantification of CD4^+^Thy1.2^+^CTV^-^ B5 T cells. **(C)** Plots and summary bar graph show MCMV-B5-specific effector and memory T cell numbers. **(D)** Plot and summary bar graph show the number within each MCMV-B5-specific Tmem subset (Tcm, TemE and TemL) as defined by CD27 and CD62L **(E)** Plot and histogram show DC and CD8α DC population gating and quantification forCD8α^-^ DC, CD8α^+^ DC and **(F)** serum IL-12p40 levels. Data shown are representative 5 mice per group and mean shown with error bars representing SEM. The Student *t* test was used; *, *P* < 0.05.

**Fig 7.**
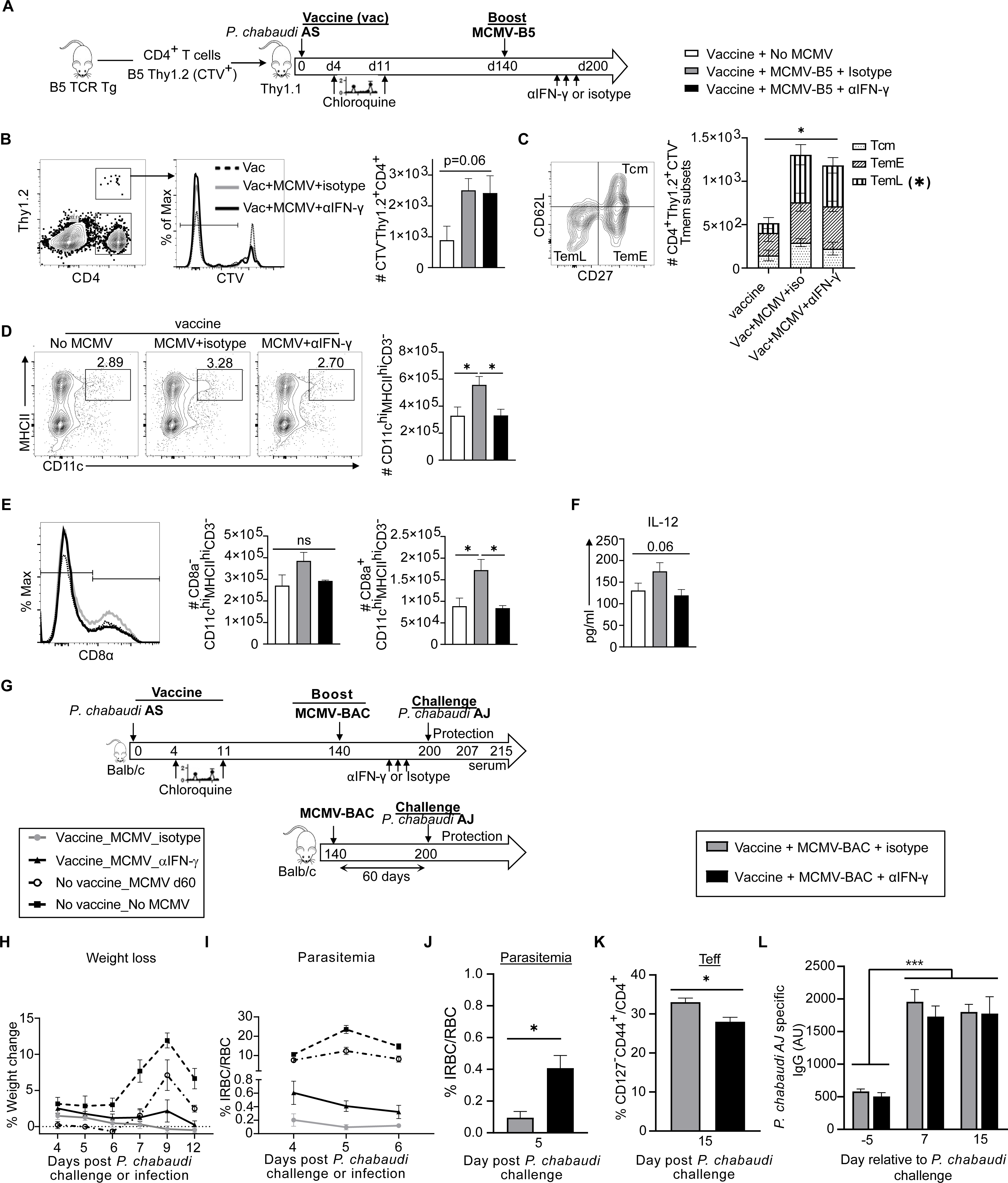
IFN-γ induced by MCMV vector drives increase in CD8α^+^ dendritic cells, and boost of long-term protection by live malaria vaccine. **(A)** Experimental schematic shows B5 TCR Tg CD4 T cells labelled with cell trace violet (CTV) were adoptively transferred into Thy1.1 mice that were then vaccinated with live malaria vaccine. Recipient vaccinated mice received MCMV-B5 boosters at day 140 and then received anti-IFN-γ antibody or isotype at indicated days (194, 196, 198) before spleen cell recovery for phenotyping at day 200 p.v. **(B)** Plot, histogram and graph show gating and quantification of B5-epitope-specific T cells (CTV^-^Thy1.2^+^) recovered on day 200. **(C)** Plot and summary bar graph show the number within each vaccine and booster-B5-specific Tmem subset (Tcm, TemE and TemL) as defined by CD27 and CD62L. * indicates significant difference between TemL subsets recovered between groups. **(D)** Plots and graphs show gating and number of total DC population (CD3^-^ CD11c^hi^MHCII^hi^), and **(E)** CD8α^-^ or CD8α^+^ DC. Plots are concatenated from 5 animals per group. **(F)** Graph of serum IL-12p40 level at day 200 p.v. before and after boosting. **(G)** Experimental schematic for infection and drug-cure vaccine combined with MCMV boosting strategy for protection to heterologous rechallenge. Age-matched mice were infected with *P. chabaudi* AS and treated with two subsequent non-curative doses of chloroquine (CQ). Then they received MCMV booster at day 140. Some mice received anti-IFN-γ antibody or isotype at indicated days (194, 196, 198) after *P. chabaudi* vaccination. All mice were reinfected with *P. chabaudi* AJ strain at 200 days p.v. Graphs showing **(H)** weight loss and **(I)** parasitemia after *P. chabaudi* AJ challenge or infection. Comparisons of vaccinated or non-vaccinated groups are significantly different by ANOVA and Tukey’s multiple comparisons significance is not shown for the sake of simplicity (*p* < 0.01, for all time points for parasitemia and at day 9 p.c. for weight loss). **(J)** Graphs showing parasitemia on day 5 *P. chabaudi* AJ challenge after IFN-γ neutralization (black bar) compared to isotype (grey bar). **(K)** Graph of fraction of polyclonal effector T cells (CD127^-^) responding to challenge (d15 p.c.). **(L) graph of** serum level of *P. chabaudi* AJ-specific IgG antibody at indicated days (-5, 7 and 15) relative to challenge. Data shown represent 5 animals per group. Mean shown with error bars representing SEM. Groups were analyzed using ANOVA followed by Tukey’s post-test for all except (J and K) which were analyzed with Student *t* test. *, *P* < 0.05; **, *P* < 0.01, ***, *P* < 0.001; n.s., not significant.

### IFN-γ generated by the MCMV booster drives prolonged protection of live malaria vaccine

Given the ability of MCMV to prolong protection from challenge and to promote IL-12 in an IFN-γ dependent manner, we tested mechanisms of the prolonged protection. Mice were vaccinated and then boosted with MCMV at day 140, as in **Fig 4**, with adoptive transfer of B5 T cells to determine the effect on vector-driven T cells. B5 T cell recipients were immunized with live malaria vaccine and boosted with MCMV-B5 at day 140 p.v., as illustrated in **Fig 7A.** Half of these animals received neutralizing antibody to IFN-γ, or isotype control antibody, days 194-198 p.v. (days 54-58 after the MCMV-B5 booster). T cell and antigen-presenting cell phenotypes were assessed at day 200 after malaria immunization, in the absence of challenge. Live *P. chabaudi* vaccination plus MCMV-B5 booster tended to increase survival of B5 T cells compared to live vaccine alone, as above, though this effect did not reach significance in this experiment (**Fig 7B**). Neutralization of IFN-γ did not have an effect on specific malaria vaccine-driven, MCMV-B5 boosted B5 T cell numbers, or the composition of Tmem subsets (**Fig 7C**). These data indicate that IFN-γ does not promote T cell survival or Tem differentiation directly, after MCMV-B5 boost, supporting the specific effect of MCMV-B5 epitope on the specific T cell responses. Staining for innate cell types, CD11c^hi^ dendritic cells (DC) were found to be increased by MCMV boost (**Fig 7D**). Furthermore, *in vivo* neutralization of IFN-γ significantly reduced the DC numbers compared to isotype control (**Fig 7D**). This innate effect was seen most strikingly in the CD8α^+^ DC (CD3^-^ CD11c^hi^MHCII^hi^CD8α^+^) population (**Fig 7E**). We also assessed NK cells, and monocyte derived subsets including monocytes, macrophages and inflammatory monocytes to see if their numbers or activation marker expression were affected by neutralization of IFN-γ, but did not see other significant differences **(S6 Fig)**. Blocking IFN-γ after vaccination and the MCMV booster caused a trend towards a reduced serum level of IL-12p40, though this did not reach significance, as it did in MCMV infection alone on day 50 (**Fig 7F**). To test the role of MCMV booster-induced IFN-γ in prolonging vaccine-induced protection, mice were vaccinated and then boosted with MCMV-BAC at day 140, as above in **Fig 3**, and as shown schematically in **Fig 7G**. The control groups either had no live vaccine and no MCMV or no vaccine and then MCMV only. Prior to heterologous challenge with *P. chabaudi* AJ at day 200, vaccinated mice with MCMV boost received IFN-γ neutralizing antibody, or isotype control, at days 194-198 p.v. (54-58 days post-MCMV booster). MCMV boosting live vaccination showed significant protection against weight loss (**Fig 7H)** and parasitemia (**Fig 7I)** compared to the adjuvanting effect of MCMV or unvaccinated control mice. Blocking IFN-γ before challenge significantly reduced the booster effect of the MCMV-BAC on protection at day 200 (**Fig 7J**). This was measured as a significant increase in parasitemia in the IFN-γ neutralized animals (0.4%), compared to the isotype group (0.09%), on day 5 post-challenge. While this is clearly a non-specific effect, as there is no B5 epitope here, polyclonal T cell phenotyping on day 15 post-challenge again showed that blocking IFN-γ before reinfection significantly reduced the fraction of Teff generated during *P. chabaudi* challenge, and was associated with higher parasitemia, suggesting a role for antigen presentation during challenge that was not tested here (**Fig 7K**). Serum levels of *P. chabaudi* AJ-specific IgG increased after challenge, but there was no significant difference from neutralization of IFN-γ (**Fig 7L**). Collectively, these data suggest a novel innate pathway where viral vector-induced IFN-γ drives an increase in CD8α+ DCs, which promote increased protective IL-12 and Teff upon challenge. We have identified two novel mechanisms of adjuvanting supporting the promise of CMV-based vaccines, and defined a promising line of research for prolonging protection from *Plasmodium* infection. Testing the full capacity of this strategy would require the identification of more protective T cell epitopes.

## Discussion

Alternative high-efficacy approaches for all malaria vaccine continue to be developed, but little engineering has been focused on increasing longevity. These data show significant promise for use of chronic vaccination to prolong the current short half-life of vaccines against systemic infections requiring CD4 T cell memory. In summary, we demonstrated an adjuvant effect dependent on the MCMV vector-induced IFN-γ that enhances CD8α^+^ dendritic cell numbers, and IL-12 production upon challenge. In addition, we demonstrate two basic mechanisms for the extension of protection provided by the chronic MCMV booster to a typical short-lived malaria vaccine. The MCMV vaccine-expressing a malaria epitope prolongs survival of epitope-specific CD4 Teff and Tem. Polyclonal effector T cell numbers also increase when challenge occurs in the presence of MCMV-induced IFN-γ, suggesting that the underlying mechanism of protection from challenge is an increased Th1 response to *Plasmodium* due to increased IL-12 downstream of higher dendritic cell numbers at the time of challenge.

We observed a strong adjuvant effect of the MCMV vector itself on *P. chabaudi* infection. The effect of MCMV alone reduced *P. chabaudi* challenge parasitemia by about one half. The adjuvant effect alone also drove a significant fraction of the prolonged protection seen from using the MCMV vector in a boosting strategy, as demonstrated by neutralizing the IFN-γ induced by MCMV-BAC before challenge. MCMV and gamma herpesvirus 68 also have an adjuvant effect against influenza virus and *Listeria monocytogenes*, but not West Nile virus [22, 23], indicating some specificity. This specificity may be due to latency. Gamma Herpesvirus 68 with a point mutation in ORF73, a gene required for latency, showed reduced protection from *Listeria*, and also reduced IFN-γ levels and chronic macrophage activation. HSV-1 and Sindbis viruses, which do not establish latency, did not have this protective effect [22]. MCMV-induced cross-protection against influenza also lasts through early and established (5-12 weeks), but not long-standing (9 months) MCMV latency. Production of IFN-γ during the latent phase of infection has also been implicated in MCMV-induced protection to *Listeria* and *Yersinia* [22], influenza; which was abrogated in IFN-γ deficient mice [23]. Our work extends these findings mechanistically by showing that neutralizing IFN-γ late in MCMV infection reduced the number of CD8α^+^ dendritic cells in the spleen, and reduced the amount of some cytokines and chemokines IL-12, CXCL-9 and CXCL-10 and the number of CD4 Teff, but increased IL-10 cytokine level on challenge. While we did not specifically test for latency, our vectors showed similar kinetics and outcomes to previous studies suggesting infection with our vectors would be latent in our mouse model by day 40 [24]. Mechanisms of this protection had not been specifically established.

Decay of immunity, and failure to maintain *Plasmodium*-specific CD4 T cells make the development of an effective and long-lived malaria vaccine challenging. Both natural immunity to the parasite, and vaccines, generate immunity that decays due to a progressive loss of malaria-specific T cells limiting long-lived protection [3, 4, 30, 31]. To our knowledge, there are no current vaccine formulations that drive long-lived CD4 T cell responses [2, 32]. Therefore, the potential of CMV to induce a prolonged T cell response with the desired phenotype was used to extend vaccine protection. In this study, a two-step vaccine strategy that first induces a long-lived protective antibody response, and then boosts with an MCMV strategy was chosen for the potential to extend effector/memory CD4 T cells responses over a longer period to prolong protection. While the protective T cell epitopes are not well defined, using a parasite derived CD4 T cell epitope allowed us to test this strategy in a pre-clinical setting. Boosting with the MCMV vector did prolong the protection generated by the live infection and-drug cure malaria vaccine as evidenced by reduced challenge parasitemia after boost. The data show that animals that received live malaria vaccination became susceptible to heterologous re-infection by day 200 p.v., while the MCMV-B5 boost maintained the protection seen at day 60 until day 200. All malaria vaccines tested in human studies thus far, including the most advanced vaccine RTS,S, show insufficient longevity of protection [25].

CMV induces a robust and long-lasting T cell response including both effector and memory components [20, 33]. Hansen et al. showed recently that RhCMV-expressing *Plasmodium knowlesi* antigens, generated a parasite antigen-specific immune response that delayed parasite growth upon sporozoite challenge via generation of CD8 T cell responses [16]. However, whether an MCMV vector could be used to boost pre-existing specificities has not been tested, and only a few studies have investigated CD4 T cell responses to MCMV [34]. Here we show that the MCMV vector can be used to specifically boost CD4 epitope-specific effector T cell responses, which is one of the primary gaps in our current malaria vaccine arsenal. Both CD4 Teff and Tmem phenotypes are generated in *P. chabaudi* infection, and both promote protection [8, 10]. Here, we observed that MCMV-B5 increased epitope-specific T cells, and specifically generated GC-Tfh and promoted maintenance of the Tem phenotype and effector functions (IFN-γ, TNF and IL-2) past day 200. CD4 T cells are crucial for protective immunity to *P. chabaudi* as they produce cytokines that both promote phagocytosis and B cell help [35]. Our previous findings revealed the power of Teff in protection from persistent *P. chabaudi* with help from B cells. Adoptive transfer of B5 effector CD4 T cells reduced the peak of *P. chabaudi* parasitemia by half [10]. While memory T cells are less effective, CD27^-^ TemL do reduce parasitemia, and are the most protective memory phenotype. In this study, we showed that the MCMV-B5 vector specifically activated and maintained both adoptively transferred epitope-specific T cells and MCMV-specific CD4^+^ T cells over 2 months. Interestingly, Teff were clearly present among the few m78-Tetramer MCMV-specific CD4 T cells over 2 months, but did not reach significant levels compared to uninfected controls. This may reflect the small number of cells detected, and so we used the B5 TCR Tg to test for MCMV-induced proliferation at late time points, however this assay had high background. Consistent with this, MCMV-B5 maintained both Teff and the potentially protective late effector memory phenotype T cells generated by *P. chabaudi* infection. While we do not present evidence here that the protection provided is due to increased survival of B5 T cells, their numbers are boosted at a timepoint when live vaccine induced T cells and protection would otherwise decay, though antibody levels remain constant. It should be noted that the B5 specificity is not especially potent in protection, as the effect of this one specificity is only seen in immunocompromised animals. CD8 T cells specific for some MCMV loci are given to inflationary activation, even after 15 weeks of MCMV infection [36]. The locus where B5 is expressed, IE1, can be a memory inflation inducing locus for CD8 T cells [37]. CD4 T cell responses to MCMV are even less well understood. Therefore, additional T cell epitope discovery in this field will aid in the long-term goal of developing durable vaccine induced protection.

The MCMV vector increased CD8α^+^ DCs and IL-12 both before challenge, and in response to the parasite. Induction of IL-12 upon *P. chabaudi* challenge was reduced in the context of prior neutralization of IFN-γ. Considering that CD8α^+^ DCs are a key source of IL-12 production in many infections [38], and also present antigen to T cells upon infection, the reduced IL-12 level in response to *P. chabaudi* challenge after IFN-γ neutralization could be directly linked to the change in Teff cell numbers seen. While IL-12 is more commonly associated with stimulating IFN-γ production, IL-12 production by DC and macrophages can also be promoted by IFN-γ present in the milieu, representing a positive feedback mechanism [39, 40]. Therefore, prolonged IFN-γ appears to be maintaining CD8α^+^ DCs, an IL-12 producing cell type, suggesting a novel mechanism for this feedback. As IL-12 deficient mice have higher *P. chabaudi* parasitemia than WT [41], or Ifng deficient animals [42], the reduced IL-12 response during challenge is likely to explain the reduced control of parasites seen upon neutralization of IFN-γ. While more research will be required to establish if the increase in dendritic cells late in MCMV infection is sufficient to drive the increased IL-12 response or the increase in Teff upon challenge, the increase of CD8α^+^ DCs supports this interpretation.

Prior work has shown that giving *P. chabaudi*-infected mice recombinant IFN-γ increased the number of polyclonal CD4 Tem, and also prolonged heterologous immunity. In our system, generation and maintenance of Tem is epitope specific, and not affected by blocking IFN-γ generated during MCMV infection. Our previous work showed that both memory T cell differentiation and prolonged infection contribute to maintenance of Tem numbers in *P. chabaudi* infection [10]. It remains to be proven whether the MCMV-BAC driven increase in DCs, and/or priming of them to respond/present antigen, leads directly to enhanced responsiveness of parasite specific T cells/expansion and clearance of parasite by Th1-promoted macrophage activation and Tfh-promoted antibody production. It is also possible that antigen presentation and Th1 response are reduced with the presence of detectable anti-IFN-γ auto-antibody [43]. Exploring this hypothesis to determine potential effects of anti-IFN-γ antibody, we found that administration of neutralizing antibody up to 2 days before infection with *P. chabaudi* does not have a significant effect on peak parasitemia, though there may be a trend towards increasing parasitemia. Indeed, in Meding et al, where IFN-γ was neutralized during the infection, the change in parasitemia was very small [42], similar to the effects of anti-IFN-γ seen on primary infection in our study. There is a clear reduction of these IFN-γ-inducible chemokines (CXCL9 and CXCL10) in the serum upon infection after anti-IFN-γ. This suggests that anti-IFN-γ directly or indirectly reduces their production in response to the parasite. An indirect example would be that these chemokines can be made by DCs, the reduction of IFN-γ induced chemokines could reflect the changes on DC number that we have documented downstream of MCMV. Our results show that although neutralizing antibody persists into the challenge phase, the largest effect of anti-IFN-γ after boosting, requires MCMV-induced IFN-γ, and is not a direct effect on parasitemia.

Neutralization of IFN-γ led to a loss of the extended protection provided by MCMV boosting and protection from *P. chabaudi* infection. Teff were reduced on challenge after IFN-γ neutralization, which is particularly striking given the increase in parasite in anti-IFN-γ pre-treated animals. IFN-γ production has been previously reported in MCMV and HCMV infection during the period coinciding with latency [23]. Data from our laboratory and others establishes that IFN-γ expression during persistent *P. chabaudi* infection, or recombinant IFN-γ also promote protection from heterologous re-infection at day 200 [12, 44]. It was also reported that rIFN-γ increased the number of polyclonal Tem at day 200 after *P. chabaudi* infection [44].

We have used basic knowledge of T cell differentiation in *P. chabaudi* infection to reverse engineer a novel chronic vaccine strategy. We demonstrate a method and a mechanism to prolong protection by a live malaria vaccine, which alone is short-lived, due to a loss of specific T cells. Our work is novel as it indicates that MCMV works as a booster for CD4 T cell responses both by maintaining specific effector and effector memory T cells and also as an adjuvant driving protective IFN-γ production. The increase in dendritic cell numbers and increased IL-12 induced by MCMV, together with and a trend towards increased Th1 cytokines upon challenge, suggest that ultimately, a stronger T cell response to parasite explains the prolonged protection. This strategy has the potential to resolve the major issue of short-lived protection generated by malaria vaccines, as well as help understand how natural immunity could be prolonged. There are safety issues to work out before CMV vectors could be used in humans. However, superinfection, the ability to re-infect CMV carriers, and a varied T cell response are two benefits of this approach [45, 46]. Further understanding of the mechanisms of this exciting new model could lead to more rapidly translatable tools in the quest to prolong immunity to persistent or recurrent pathogens without introducing another pathogen. This could be applicable to malaria, a disease that kills up to half a million people a year; as well as other persistent or recurrent pathogens.

## Materials and methods

### Ethics statement

All experiments were carried out according to protocol approved by the University of Texas Medical Branch Institutional Animal Care and Use Committee (IACUC).

### Mice, Parasites and Immunization

All mice used in our study were maintained in the UTMB animal facility under specific pathogen-free conditions with access to food and water *ad libitum.* Recipient Thy1.1 BALB/cByJ were backcrossed to BALB/cJ (N4; The Jackson Laboratory, Bar Harbor, ME). B5 T cell receptor transgenic (TCR Tg), which recognizes a peptide from merozoite surface protein-1 (MSP-1; 1157-1171, ISVLKSRLLKRKKYI/I-E^d^) was backcrossed to BALB/cJ (N10). B5 TCR Tg mice, and *P. chabaudi chabaudi* AS stocks, were a kind gift from J. Langhorne (Francis Crick Institute, London, UK). *P. chabaudi* AJ (MRA-740, a cloned line that is highly related but heterologous to AS [47, 48] was purchased from BEI Resources, formerly MR4. B5 TCR Tg mice were typed using primers Vα2, 5′-gaacgttccagattccatgg-3′ and 5′-atggacaagatcctgacagcatcg-3′, and vβ8.1, 5’ cagagaccctcaggcggctgctcagg-3′ and 5′-atgggctccaggctgttctttgtggttttgattc-3′ [8]. Both male and female recipients were used, however no sex differences in infection were noted. All adoptive transfer experiments were done using age- and sex-matched recipient and donors.

For infection and drug cure vaccination, the protocol was performed as previously documented in mice and humans [3, 27]. Six-to nine-week old mice were infected (10^6^ *P. chabaudi)* and treated with two subsequent noncurative doses of chloroquine (10 mg/kg body weight/day; Sigma-Aldrich), when parasitemia reached 1-2% of erythrocytes. Parasites were counted by light microscopy in thin blood smears stained with Giemsa (Sigma-Aldrich, St. Louis, MO). Parasitemia was monitored by microscopic examination of thin blood smears from the tail. Thin blood smears were stained with Giemsa (Ricca chemical company, Arlington, TX), and parasites were counted by light microscopy.

### Construction of MCMV and viral infection

A vaccine vector, MCMV-K181-B5, was generated as follows. MCMV-K181-Perth was genetically modified to carry the MHC II-E^d^-restricted B5 epitope (ISVLKSRLLKRKKYI) from the *Plasmodium chabaudi* Merozoite Surface Protein MSP1 (1157–1171) in frame with the C-terminus end of the MCMV IE1 gene (**Fig S1 A**). The B5 epitope was inserted into the MCMV-K181 genome using bacterial artificial chromosome recombineering, as described previously [49]. The resulting virus was denoted MCMV-B5. Mice were infected by injecting 5x10^3^ PFU of salivary gland-propagated MCMV-B5 virus or the parental virus, MCMV-K181-Perth (MCMV-BAC). Viruses were injected i.p. in 0.2ml of PBS.

### Virus quantification

MCMV-B5 or MCMV-BAC viral copy number of CMV/β-actin ratios were determined in different organs in infected BALB/cJ mice. DNA was purified from tissue (liver, lung, and salivary gland) using the Gentra Puregene Kit (Qiagen, Valencia, CA) following manufacturer’s instructions. All tissues were stored at -80°C in PBS for DNA, or Trizol (Thermo Fisher Scientific, Carlsbad, CA) for RNA purification, until sample processing. qPCR was performed with 50 ng of purified DNA. Specific primers for MCMV immediate early (IE1) gene (NCBI Accession No. M11788) 5-tca gcc atc aac tct gct acc aac-3 and 5-gtg cta gat tgt atc tgg tgc tcc tc-3 and for murine β-actin (Accession No. M12481) 5-gct gta ttc ccc tcc atc gtg -3 and 5-cac ggt tgg cct tag ggt tca-3. Serial dilutions of a control plasmid were used to generate a standard curve for both MCMV-IE1 and β-actin, as described previously, and qPCR of viral DNA has been shown to correlate quantitatively with the plaque assay throughout the viral infection [50]. All sample measurements were performed in triplicate. Real time PCR (VIIA, AB Biosystems, Life Technologies, Carlsbad, CA).

### Cell culture with serum, RNA isolation and real-time PCR analysis

RAW264.7 macrophage-like cell line was cultured in RPMI medium (Gibco) supplemented with 10% (v/v) fetal bovine serum and antibiotics (100 U/ml penicillin and 100 lg/ml streptomycin), Sodium Pyruvate (100x, Gibco), 45% D-Glucose (100x, Sigma), L-glutamine (100x, Corning) at atmosphere of 5% CO2. RAW264.7 cells were incubated with sera form IFN-γ neutralizing mice with or without recombinant IFN-γ for 24h. Total RNA was extracted with Direct-Zol RNA reagents (Zymo research, R2025) as described by the manufacturer. cDNA was synthesized from total RNA using the iScript cDNA Synthesis kit (Bio-Rad). Real-time PCR was performed with cDNA using the SYBR Green Supermix (Bio-Rad). The β-actin gene was used for normalization. Relative changes in gene expression were calculated using the comparative threshold cycle (Ct) method. The following primers were used: CXCL-10 5’-cctgctgggtctgagtggga-3’ and 5’-gataggctcgcagggatgat-3’; β-actin 5-gctgtattcccctccatcgtg-3 and 5- cacggttggccttagggttca-3.

### Adoptive transfers and *in vivo* studies

MACS-purified CD4^+^ T cells from young, uninfected B5 TCR-Tg mouse spleens (Miltenyi San Diego, CA) were labelled with cell trace violet (CTV) and adoptively transferred (2×10^6^) into Thy1.1 congenic mice. For cytokine neutralization, three doses of anti-IFN-γ antibody (H22, 0.5 mg) or isotype control antibody (PIP, 0.5 mg); or a combination of anti-IL-12p40 (C17.8, rat IgG2a, 300 μg) and anti-IL-18 antibody (YIGIF74–1G7, control Rat IgG2a, 200 μg) were given, every other day. All *in vivo* antibodies were from BioXCell (Lebanon, NH).

### Flow cytometry

Single-cell suspensions from spleens were prepared at the indicated time points after infection in HBSS) with HEPES (Thermo Fisher Scientific, Waltham, MA) using gentleMACS (Miltenyi, San Diego, CA) and incubated in RBC lysis buffer (eBioscience, San Diego, CA). Single cells were stained extracellular in PBS, 2% FBS, and 0.1% sodium azide with antibodies. For T cells: anti-CD127 (A7R34), anti-PD-1 (RMP1-30), anti-CD62L (MEL-14), CD4 (RM4-5), (anti-CD90.2 [30-H12], anti-CD44 [IM7], and anti-CD27 [LG-7F9]) (all from eBioscience/Invitrogen and Biolegend, San Diego, CA). Phenotype of polyclonal CD4^+^ T cell subsets and *Plasmodium* B5-specific CD4^+^ T cell subsets were identified with gating strategy as schematically depicted in **S7 Fig**. For follicular T cell staining with CXCR5, cells were stained in PBS + 0.5% BSA + 0.1% sodium azide + 2% Normal Mouse Serum (NMS) and 2% FBS (Sigma, St. Louis, MO). Rat anti-mouse purified CXCR5 (2G8, BDbioscience, San Jose, CA, 1 hr., 4°C) was followed by biotin-conjugated AffiniPure Goat anti-rat (H+L, Jackson ImmunoResearch, West Grove, PA, 30 min, 4°C) followed by Streptavidin-eFluor 450. For antigen presenting cells were stained with anti-Ly6C, anti-Ly6G (1A8), anti CD11b (M1/70), anti-CD11c (N418), DX5, MHCII/I-A/Eb (M5/114.15.2). For intracellular cytokine staining, cells were cultured for 5 hours at 37°C, 5% CO_2_ in the presence of GolgiPlug (BD Bioscience) before extracellular staining and then fixed with 2% paraformaldehyde overnight. Cells were then permeabilized using BD 10X Permeability buffer diluted with water, and stained with anti-IFN-γ (XMG1.2), anti-TNF (MP6-XT22), anti-IL-2 (JES6-5H4). Virus-specific CD4^+^ T cells were identified using m78/I-A^d^ tetramers, with peptide (SQQKMTSLPMSVFYS) and CLIP/I-A^d^ as tetramer control (PVSKMRMATPLLMQA) and CD8^+^ T cells with H2D^d^ restricted m45-tetramer peptide 507-VGPALGRGL-515 [20, 21] (both from NIH Tetramer Core Facility, Emory, Atlanta, GA). Tetramer staining was performed within 6 months of receiving the reagents at 4C for 1 hour before/after surface and intracellular staining. Cells were acquired on a LSR II Fortessa using FACSDiva software (BD Biosciences, San Jose, CA). Compensation was performed in FlowJo using single-stained splenocytes (with CD4 in all colors). All data analysis was performed using the FlowJo software package (version 10, FlowJo, BD San Jose, CA). Cell numbers were calculated as the % of recovered Tg cells/lymphocytes x the number of lymphocytes, except for **Fig 6** and **Fig 7** which the recovered cells were counted by inclusion of counting beads (Accu-Check, Molecular Probes, Thermo Fisher Scientific) in the FACS sample according to manufacturer’s instructions and calculated using the equation: Total cell number = ((all cell event count / bead event count) x bead conc. (per µl) x (bead volume / cell volume)) x volume of original sample. This formula only works if the whole volume of FACS tube is collected.

### Cytokines and antibody measurements

Cytokines and chemokines (CXCL10 (IP-10), IFN-γ, IFN-β, IFN-α, IL-10, IL-12p40, IL-15, IL-18, IL-6, and TNF-α) were measured in the plasma by Legendplex according to manufacturer’s protocol (Biolegend, San Diego, CA). Plasma samples were collected at the indicated days and were stored in -80°C until analysis. All data were acquired on an LSR II Fortessa flow cytometer using FACSDiva software, version 8 (BD Biosciences, San Jose, CA). The concentration of each analyte is determined with cytometric data based on a standard curve using the LEGENDplex™ data analysis software (Biolegend, San Diego, CA). *P. chabaudi* AJ-specific IgG was evaluated in plasma at indicated days by ELISA, as described previously [5]. Plates were coated with *P. chabaudi* AJ parasite lysate overnight at 4°C in PBS with 0.05% Sodium Azide. Plates were blocked with 1% BSA, 0.3% Tween 20, 0.05% Sodium Azide and then sera were incubated. Bound antibody was detected using Alkaline Phosphatase-conjugated goat anti-mouse IgG (Southern Biotech, Birmingham, AL). The reaction was revealed with a p-Nitrophenyl phosphate disodium salt (Sigma, USA) solution (1mg/ml) in diethanolamine buffer. Plates were analyzed with an Omega plate reader (FLUOstar Omega BMG LABTECH Inc, Durham, NC, USA).

### Statistical analysis

All data were analyzed by ANOVA, followed by Tukey’s or two-tailed Mann-Whitney test or two-tailed unpaired Student t test, where indicated. All data are presented as mean ± SEM. Statistical tests were performed using the statistical software package in Prism (version 9.2, GraphPad Software, USA): *p < 0.05, **p ≤ 0.01, ***p ≤ 0.001 and n.s., not significant.

### Supporting information

## Acknowledgments

We would like to thank Florentin Aussenac, Victor H. Carpio, Nadia D. Domingo, Lucinda Puebla-Clark and Peter Fleming for feedback and discussion. Thanks to Maria I. Giraldo Giraldo and Gregg Milligan for technical advice. We thank the NIH Tetramer Core Facility at Emory, Atlanta, GA for generating both the m78-Tetramer and m-45-Tetramer used in this study. This work was supported by the James W. McLaughlin Fellowship Fund (KG) and NIH R01AI135061, R01AI089953 (RS, KG).

## Author Contributions

### Conceptualization

KG, MGB, MAD, RS

### Investigation

KG, SAI

### Formal Analysis

KG, SAI, RS

### Methodology and Resources

KG, SAI, MLO, MAD, MGB

### Funding acquisition

KG, RS, MGB, MAD

### Project Administration, Supervision

RS, MAD

### Writing – Original Draft Preparation

KG

### Writing – Review & Editing

RS, MGB, MAD

---

S1 **Fig. *Plasmodium chabaudi* MSP-1 B5 antigen-expressing MCMV vaccine vector.**

**(A)** *P. chabaudi* merozoite surface protein MSP-1 (B5:1157–1171) B5 epitope is expressed as an MCMV immediate early gene. B5 epitope was introduced into an MCMV-K181 bacterial artificial chromosome (BAC) using recombineering. **(B)** The indicated organs were collected from MCMV infected mice at different times post-MCMV infection, homogenized, and MCMV titres were determined by plaque assay. The dashed line represents the limit of detection (L.O.D.) for each organ. Mean ± SEM are plotted with three mice for each time point. **(C)** Time course of qPCR assessment of MCMV infection in multiple organs. Mice were infected with MCMV-B5 or MCMV-BAC and organs were collected at indicated times for vector quantification with n=6 per time point. Copy number of CMV/β-actin (±SEM) ratio and L.O.D. of the assay are shown. L.O.D was calculated using data from uninfected animal. (**D**) Mice were infected with MCMV-BAC or MCMV-B5 and splenocytes were recovered 15 days later for MCMV-induced polyclonal T cell phenotype. Plots show gating and graphs show numbers of polyclonal CD4^+^ T cells effector (CD127^-^) and memory (CD44^hi^CD127^hi^) phenotype cells recovered after MCMV-B5 compared to MCMV-BAC. **(E)** Enumeration of MCMV-specific m78-Tetramer-positive CD4 T cells upon MCMV-B5 compared to MCMV-BAC from day 15 MCMV-infected mice. Plots represents control CLIP tetramer or m-78-tetramer staining after MCMV-B5. Graph shows number of MCMV-specific m78 Tet+ T cells (±SEM). **(F)** Enumeration of MCMV-specific m45-Tetramer-positive CD8 T cells. Mice were infected with MCMV-B5 and splenocytes were recovered 11 and 60 days later for CD8 T cell quantification by staining with m45 tetramers. Plots and graph show m45-Tet^+^ CD8^+^ T cells from uninfected and MCMV-B5 infected mice gating on vector specific T cells (+/- SEM). Data shown are representative of two experiments with 3 or 5 mice each. Groups were analyzed using One-Way ANOVA followed by Tukey’s post-test with *, *P* < 0.05; **, *P* < 0.01.

**S2 Fig. Persistent MCMV-B5 infection induces vector-specific T cell response and may promote continued B5 TCR Tg T cell proliferation.**

**(A)** Experimental schematic shows mice were infected with MCMV-B5, and splenocytes were recovered 11 or 60 days later to quantify MCMV m-78-specific Tetramer^+^ CD4 T cells. (**B**) Plots and graph show uninfected and m78 tetramer-specific T cell numbers. Virus-specific CD4^+^ T cells were identified using m78/I-A^d^ tetramers, with peptide (SQQKMTSLPMSVFYS) and CLIP/I-A^d^ as tetramer control (PVSKMRMATPLLMQA). (**C**) Plots and graphs show the number of MCMV-specific-Teff (CD44^+^CD127^-^) and Tmem (CD44^hi^CD127^hi^) in the spleen gated on MCMV m78-tetramer CD4^+^ T cell at day 11 and 60 p.i.. Data shown are representative of two experiments with 5 mice each and error bars representing SEM. **(D)** Experimental Schematic shows MCMV-BAC or MCMV-B5 infection of different groups of age-matched Thy1.1 mice at indicated times (0, 30 and 50 days) followed by adoptive transfer of CTV^+^ B5 TCR Tg CD4 T cells to all groups 5 days before phenotyping, and 55 days after first infection. **(E)** Plots and graphs showing gating and frequency of CTV-B5 T cells out of CD4 T cells, or number of proliferated B5 epitope-specific CD4 T cells in response to MCMV-BAC or MCMV-B5 infection at different time points. Data are representative of three independent experiments with 5 animals per group. Mean shown with error bars representing SEM. Groups were analyzed using One-Way ANOVA followed by Tukey’s post-test with *, *P* < 0.05; **, *P* < 0.01; ***, *P* < 0.001.

**S3 Fig. Decay of *P. chabaudi* protection by days 200**

**(A)** Experimental schematic shows age-matched mice were infected with *P. chabaudi* AS for assessment of protection to heterologous challenge with *P. chabaudi* AJ 60 or 200 days later. Graphs show **(B)** the percentage of parasitemia (iRBC/RBC) after heterologous challenge, and **(C)** *P. chabaudi* AJ-specific IgG antibody serum level at day 7 post-challenge. Data are representative of two independent experiments with five mice per group showing mean, with error bars representing SEM. Groups were analyzed using One-Way ANOVA followed by Tukey’s post-test with *, *P* < 0.05; **, *P* < 0.01; ***, *P* < 0.001.

**S4 Fig: Evaluation of neutralizing effect of anti-IFN-γ antibody**

**(A)** Experimental schematic shows mice were infected or not infected with MCMV-BAC to assess the neutralizing effect of anti-IFN-γ naïve mice or in MCMV-induced protection to *P. chabaudi*. Blocking of IFN-γ *in vivo* with anti-IFN-γ antibody or isotype control antibody was performed at days 34, 36 and 38 prior to the challenge or infection with *P. chabaudi* at day 40 in both MCMV infected and naïve mice. Graphs show percentage of parasitemia in **(B)** no MCMV mice and **(C)** MCMV vaccinated mice. Relative CXCL-10 mRNA level normalized with β-actin from Raw 264.7 cells cultured for 24h with sera from: (**D**) 2 days after anti-IFN-γ administration, (**E**) day 7 post *P. chabaudi* challenge, (**F**) 2 days after anti-IFN-γ administration in presence of rIFN-γ (25ng/ml), and (**G**) day 7 post *P. chabaudi* challenge administration in presence of rIFN-γ (25ng/ml). Data shown represent 5 animals per group and mean shown with error bars representing SEM. Groups were analyzed using Student *t* test (C, D and F) or One-Way ANOVA (B, E and G) with *, *P* < 0.05; **, *P* < 0.01; ***, *P* < 0.001.

**S5 Fig. Combined IL-12 and IL-18 neutralization did not affect protection induced by MCMV.**

**(A)** Experimental schematic shows mice infected with MCMV-BAC and the combination of anti-IL-12p40 and anti-IL-18 antibody or isotypes were given post MCMV infection. Graphs show percentage of parasitemia **(B)**, weight loss **(C)**, and hypothermia **(D)** upon combined IL-12 and IL-18 neutralization. Data shown represent 5 animals per group and mean shown with error bars representing SEM and analyzed using multiple comparisons Student *t* test; n.s., not significant.

**S6 Fig. IFN-γ induced by MCMV vector does not change most antigen-presenting cells types and their activation significantly.**

**(A)** Experimental schematic shows mice infected with MCMV-B5 and received anti-IFN-γ antibody or isotype at indicated time points. Splenic antigen-presenting cells types (monocytes, macrophages and inflammatory monocytes) and NK cells were phenotyped at day 50 p.i.. **(B)** Plots and graphs show gating and quantification of CD3^-^CD11b^-^Ly6C^+^, CD3^-^ CD11b^+^Ly6C^+^, CD3^-^ CD11b^+^Ly6C^-^ cells. (C) Histograms and graphs show MFI of CD3-CD11b+Ly6C-expressing activation marker MHCII. (D) Histograms and graphs show MFI of CD3-CD11b+Ly6C-expressing activation marker CD86. (**E**) Plots and graphs show gating and quantification of CD3^-^ DX5^+^MHCII^-^ NK cells.

(**F**) Experimental schematic shows Thy1.1 mice vaccinated with live malaria vaccine. Then vaccinated mice received MCMV-B5 boosters at day 140 and then received anti-IFN-γ antibody or isotype at indicated days (194, 196, 198) before spleen cell recovery for cells phenotyping at day 200 p.v. **(G)** Plots and graphs show gating and quantification of CD3^-^ CD11b^-^Ly6C^+^, CD3^-^ CD11b^+^Ly6C^+^, CD3^-^ CD11b^+^Ly6C^-^ cells. **(H)** Histograms and graphs show MFI of macrophages (CD3^-^CD11b^+^Ly6C^-^) expressing activation marker MHCII. **(I)** Histograms and graphs show MFI of inflammatory monocytes (CD3^-^ CD11b^+^Ly6C^+^) expressing activation marker MHCII. (**J**) Plots and graphs show gating and quantification of CD3^-^DX5^+^MHCII^-^ NK cells. Data shown are representative 5 mice per group. Plots concatenated per group and mean shown with error bars representing SEM. The Student *t* test or One-Way ANOVA were used.

**S7 Fig. An example of a gating strategy used to identify polyclonal CD4^+^ T cells subsets and B5-specific CD4^+^ T cells subsets**

**(Top row)** Plots show gating strategy for polyclonal CD4^+^ T cell subsets Teff (CD4^+^CD127**^-^**CD44^hi^), Tmem (CD4^+^CD127^hi^CD44^hi^); and memory T cell subsets gated on CD44^hi^CD127^hi^ as Tcm (CD62L^+^CD27^+^), Tem^Early^ (TemE, CD62L^lo^CD27^+^) or Tem^Late^ (TemL, CD62L^lo^CD27^-^). **(Bottom row)** Plots show gating strategy for B5-specific T cells: Teff (CD4^+^Thy1.2^+^CTV^-^, CD127**^-^**CD44^hi^), Tmem (CD4^+^Thy1.2^+^CTV^-^, CD127^hi^CD44^hi^) and B5 memory T cell subsets gated on CD44^hi^CD127^hi^, Tcm (CD62L^+^CD27^+^), Tem^Early^ (TemE, CD62L^lo^CD27^+^) or Tem^Late^ (TemL, CD62L^lo^CD27^-^).

